# Epitopes- Based Vaccine Design Against Foot and Mouth Disease *SAT2* Serotype from Sudanese Isolate by Using Immunoinformatic Approaches

**DOI:** 10.1101/2022.12.23.520716

**Authors:** Inas. I. Habiballa, Y. A Raouf, M Alhaj, Sana. I. Mohamed, Reham M. Elhassan, Oubi. O. Salim, Essra Mustafa, A. Mohamed, K Ahmed, M Mona, Mohamed. M. Sirdar, Mohammed A. Hassan

**Author notes:** **Corresponding author:** Inas Habiballa.

## Abstract

Foot and mouth disease (FMD) has been endemic in Sudan for decades and causes continuous outbreaks that have a direct negative impact on the animal population and prevent the exportation of animals from the country. The high diversity of FMD serotypes, especially SAT2 and A serotypes, hinders the development of effective vaccines since the most important component of vaccination is the degree of cross-protection provided by the vaccine against currently circulating field viruses. An immunoinformatic approach was utilized to predict a multi-epitope peptide vaccine design against the *SAT2* serotype from a Sudanese isolate targeting virus capsid region *P1*. The virus capsid region *P1* comprises the major immunogenic epitopes that confer protection against the FMD virus. Two predicted T-cell epitopes were identified that showed high binding affinity with MHC1 alleles (***VQRSRQSTL** and **YHAEWDTGL***) and high conservation with SAT2 African serotypes and were located within the VP1 and VP3 proteins, respectively. Only one epitope was predicted for B cells (***LPATPEDAAH***), which scored above the threshold in Bepipred linear epitope, Emini surface accessibility, and Kolaskar and Tongaonkar antigenicity and is located in VP3 protein. Molecular docking of the peptides (***VQRSRQSTL** and **YHAEWDTGL***) with the MHC1 allele showed satisfactory interaction with the binding sites of BoLA-HD6 using UCSF chimera 1.13.1 software. The peptide **VQRSRQSTL** showed remarkable hydrophobic interaction with the BoLA-HD6 allele, which was superior to the other peptides. This study is the first to propose a peptide vaccine against *FMD SAT2* serotypes from a Sudanese isolate.

## Introduction

Foot and mouth disease (FMD) virus belongs to the *Picornaviridae* family, a member of the genus *Apthovirus*. FMD is an acute and highly contagious febrile disease affecting all ruminants and cloven-footed animals (King *et al* 2011). The FMDV particle is roughly spherical in shape and about 25–30 nm in diameter. It consists of the RNA genome surrounded by a protein shell or capsid. The capsid is composed of 60 copies of the capsomers. Each capsomer consists of four structural polypeptides, VP1, VP2, VP3, VP4 and nine nonstructural proteins are cleaved by viral proteases (Rueckert 1996). The VP1, VP2 and VP3 are exposed on the surface of the virus while VP4 is located internally (Jamal and Belsham 2013). FMDV exists as seven immunologically distinct serotypes O, A, C, Asia 1, Southern African Territories; SAT1, SAT2 and SAT3 (Alexandersen and Mowat 2005). The disease causes huge economic losses to livestock industries worldwide; even though fatal cases are usually restricted to young animals and certain FMDV strains (Alexandersen *et al* 2003). FMD infected animals significantly reduce productivity and affects international trade of animals and animal-derived products. Livestock trade is severely restricted between countries with different sanitary status of the disease (Knight-Jones and Rushton 2013). FMD was firstly reported in the Sudan in 1903; the country is endemic by the disease that causes severe negative economic impact in the dairy sector and in international trade of animals (Abu ElZein 1983). Recent update of the disease situation in the country indicate the maintained activity of three serotypes; O, A and SAT2 (Raouf *et al., 2010*; Habiela *et al* 2010; Raouf *et al* 2014, 2016). Vaccination has been successfully applied as the main control measure for FMD in different South American countries, sub Saharan African countries and many parts in the world (Hunter 1998). In most cases, vaccination is formulated by means of polyvalent oil vaccines or aqueous vaccine using inactivated virus from strains previously detected in the region (Saraiva and Darsie 2004). Conventional FMD vaccines can preclude clinical infection, but like other killed antigens, they do not induce broadly reactive long-term protection. They require multiple vaccinations to preserve adequate levels of individual and herd immunity, and need periodic inclusion of new viral strains into the vaccine formulation to cover different virus topotypes that may not be covered by the existing vaccines. Other important shortcomings of inactivated vaccines include i) short shelf life, ii) the need for adequate cold chain of formulated vaccines, and iii) difficulties of certain serotypes and subtypes to grow well in cell culture which is a requisite for vaccine production. To address these problems, basic research and novel technologies have been explored, such as computational vaccinology, which holds the promise of changing vaccine development in that it challenges the conventional approach in reducing the cost and period of production (Diaz-San Segundo et al. 2016, Vaishnav et al. 2014). There has been an increase in the use of computational approaches for vaccine development based on genomic information; this was termed “reverse vaccinology” (Bambini and Rappuoli 2009; De Groot et al. 2002; Major et al. 2011; Plotkin 2008; Serruto et al. 2009). Diverse immunoinformatics algorithms have been developed to envisage T- and B-cell immune epitopes for epitope vaccine development (Vaishnav *et al*. 2014). An epitope or antigenic determinant is a collection of short amino acid residues on an antigen for antibodies to recognize and specifically bind to successfully generate an immune response (Larsen *et al*., 2006; Peters *et al*., 2005a; Zhao *et al*., 2012; Sun *et al*., 2013). Recent immunoinformatic approaches have been extensively applied to recent FMD vaccine research. Liu *et al*. (2017) designed peptides mainly from VP1 protein, chemically synthesized them, and then tested the potency of the vaccine. A strong association was obtained between virus neutralization test titers and protection after challenging the animals. The presented study is the first investigative report to predict T and B cell epitopes against *FMD SUD/NK/28/2010 SAT2 polyprotein* from a local Sudanese isolate to generate a proposed candidate peptide-based vaccine that protects against FMDV.

## Material and method

### Virus

*FMDV serotype SAT2/NK/2010* was obtained from North Kordofan State, Shikan District in North West Sudan.

#### Propagation of Virus in Cell Culture

The virus was propagated in Baby Hamster Kidney (BHK) clone 21; first, the virus with dilution 1/10 was inoculated into the BHK monolayer and then incubated at 37 C for 1 hour. After the incubation, GMEM media with 10 % fetal bovine serum was added to the cells and incubated at 37 C for 24 hours. The development of CPE was observed, which included rounding of cells, cell detachment from the cell culture flask and complete destruction of the cells. The virus was passed up to 28 times.

### Molecular analysis

#### RNA extraction

Genomic RNA was extracted from cultured virus by a modified silica/guanidinium thiocyanate nucleic acid extraction method with the Qiagin kit (Boom et al., 1990).

#### PCR condition for P1 region

Genomic amplification of P1 region was performed by using forward primer *SAT2-P1*-1223F [*5-TGAACT ACC ACT TCA TGT ACA CAG-3*] and *SAT-2B208R* [5-*ACA GCG GCC ATG CAC GAC AG*-3] as a reverse primer (Habiela *et al* 2010 and Knowles *et al* 2016). One step PCR was performed in a final volume of 50ul including 3 μl of template RNA, 2 μl RT and Taq DNA polymerase enzymes (Qiagen), 2 μl SAT2 (forward, reverse) Primers, 10 μl of Qiagen 10x, 2 μl dNTP, then 29 μl of distilled water, were added to complete the volume to 50 μl. The PCR protocol was designed for SAT2 serotype according to (Reid *et al* 1999).

#### Sequencing of the PCR product

PCR product was purified and sequenced for both strands of *FMD P1* region in Macrogen company in China by using Sanger method. The obtained nucleotides were translated by Gene mark software http://opal.biology.gatech.edu/GeneMark/ to a protein with 278 amino acids in length.

#### Protein sequences retrieval

A total of 10 sequence of FMD virus SAT2 poly protein were retrieved in FASTA format from NCBI database (http://www.ncbi.nlm.nih.gov/protein) on 23^rd^ April 2019 These sequences retrieved are from different parts of Africa SAT2 serotypes which include Egypt, Nigeria, South Africa, Kenya and Saudi Arabia. The retrieved proteins strains and their accession numbers and area of collection were listed in (Table 1).

**Table (1):** The retrieved proteins strains and their accession numbers and area of collection were listed in

| **Accession number** | **Date of collection** | **Country** |
| --- | --- | --- |
| QBK46481 | 2012 | Nigeria |
| AAQ63182 | 2000 | Saudi Arabia |
| QBK46484 | 2012 | Nigeria |
| QBK46477 | 2012 | Nigeria |
| ARO74646 | 2014 | Egypt |
| AJW67262 | 2010 | Kenya |
| BBI28318 | 2018 | Ghana |
| BBI28317 | 2018 | Ghana |
| AFP33712 | 2012 | Egypt |

**Table (1):** List of peptides with their surface accessibility score and antigenicity score

| Start | End | Epitope | Length | Accessibility | Antigenicity |
| --- | --- | --- | --- | --- | --- |
| 23 | 30 | ATPEDAAH | 8 | 1.286 | 0.998 |
| 22 | 30 | PATPEDAAH | 9 | 1.575 | 1.006 |
| 21 | 30 | **\*LPATPEDAAH** | 10 | 1.036 | 1.03 |
| 20 | 30 | QLPATPEDAAH | 11 | 1.432 | 1.029 |
| 19 | 30 | TQLPATPEDAAH | 12 | 1.658 | 1.019 |

\* Refer to the conserved peptides that passed the Emini surface accessibility and Kolaskar and Tongaonkar antigenicity test.

#### Phylogeny and alignment

The retrieved sequences from NCBI and the FMD SAT2/NK/28/2010 Sudanese isolate were used in a phylogenetic analysis to determine the common ancestor of each strain and the conservancy using various tools in Mega 7 software (Kumar et al. 2016). The phylogenetic tree and alignment are presented in Figures (1).

The sequence of *FMD SAT2 serotype P1* poly protein was analyzed by different prediction tools from Immune Epitope Database IEDB analysis resource to predict the B- and T helper cell epitopes (http://www.iedb.org/) (Chevenet *et al* 2006, Hall 1991).

#### B- cell epitope prediction

##### Prediction of linear B-cell epitopes

Bepipred test from immune epitope database (http://tools.iedb.org/bcell/result/) was used as linear B-cell epitopes prediction tool to rule out passed Linear B- epitopes with score above the default threshold value of (0.0350) (Larsen *et al* 2006).

##### Prediction of surface accessibility

Emini surface accessibility prediction tool (IEDB) (http://tools.iedb.org/bcell/result/) was used to predict surface epitopes with the default threshold value 1.000 (Emini *et al* 1985).

##### Prediction of epitopes antigenicity

Kolaskar and Tongaonkar antigenicity method was used to determine the antigenic sites with a default threshold value of 1.033, (http://tools.iedb.org/bcell/result/) (Kolaskar and Tongaonkar 1990).

##### T cell epitope prediction

Analysis of peptide binding to MHC class I molecules (BolA T-cell epitope) was evaluated by the IEDB MHC I prediction tool at (http://tools.iedb.org/mhci/n), MHC-I peptide complex presentation to T lymphocytes undergo several steps. The attachment of cleaved peptides to MHC molecules step was predicted. Prediction methods can be achieved by Artificial Neural Network (ANN), ANN method was used (Peters and Sette 2005b). Prior to prediction, all epitope lengths were set as 9 mers, that bind to alleles at score equal or less than 500 half-maximal inhibitory concentration (IC50) is selected for further analysis (Sidney *et al* 2008).

##### Physicochemical properties

It is essential to define the physical and chemical parameters associated with the vaccine. The physicochemical properties of FMD SAT2 serotype P1 poly protein was analyzed using BioEdit sequence alignment editor software Version 7.2.6.0 and the chemical properties of epitopes where identified (Hall *et al* 1999).

##### Homology modeling

The modeling has been done by Raptor X structure prediction server. The reference sequence of *FMD SAT2 serotype* P1 poly protein (Maree *et al* 2011) was submitted on 3^rd^ January 2019. The file result is received at 3 January 2019, which containing the predicted 3D model and image as an attachment. UCSF Chimera (version 1.8 Chimera X version 0.1) was used to visualize the 3D-structures of protein and to allocate the sequential location of the proposed epitopes within the model. (Bui *et al* 2006).

##### Molecular docking analysis

Molecular docking was performed using Moe 2007; the 3D structures of the promiscuous epitopes were predicted by PEP-FOLD (Lamiable *et al* 2016, Thevenet *et al* 2012). BoLA-HD6, and BoLA-T2C were chosen as a model for molecular docking to predict the strength of association between the binding site and promiscuous epitopes. The crystal structures of the models were downloaded in a PDB format from the RCSB PDB resource. However, the selected crystal structures were in a complex form with ligands. Thus, to simplify the complex structure all water molecules, hetero groups and ligands were removed by Discovery Studio Visualizer 2.5 (Lamiable *et al* 2016). Partial charge and energy minimization were applied for ligands and targets. In terms of the identification of the binding groove, the potential binding sites in the crystal structure were recognized using the Alpha Site Finder. Finally, ten independent docking runs were carried out for each Peptide and results were retrieved as binding energies. The best poses for each epitope that displayed the lowest binding energies were visualized using UCSF chimera 1.13.1 software (1-11).

## Results

### Propagation of virus in cell culture

The virus growth was satisfactory in BHK 21 cell line culture; which included rounding of cells and finally complete CPE destruction (figure 1).

### Molecular characterization

PCR was conducted on cultured virus; the sample under test gave positive result for the P1 gene which codes the capsid proteins. Strong bands were detected in the ethidium bromide stained gel that correspond exactly to the expected DNA band size of the control positive DNA (1279 bp) (Knowles *et al* 2016). 1222 nucleotides were obtained after Sanger sequencing, the sequences chromatogram was viewed by Finch TV program, (http://www.geospiza.com/Products/finchtv.shtml) figure (1).

### Phylogeny and alignment

The phylogeny alignment revealed that *FMD SAT2 SUD/NK/28/2010* serotype based on protein sequence of (P1) poly protein coding region showed that it is more related to Egyptian isolates (EGY/24/2014 and EGY/9/2012) and Saudi Arabia (SAU/6/00) figure (1).

### B cell epitope prediction

Only one B cell epitope contained 10 amino acids was predicted and passed all the tests above the threshold, the result is summarized on table (1) and figures 2, 3 and 4.

**Figure (1).**
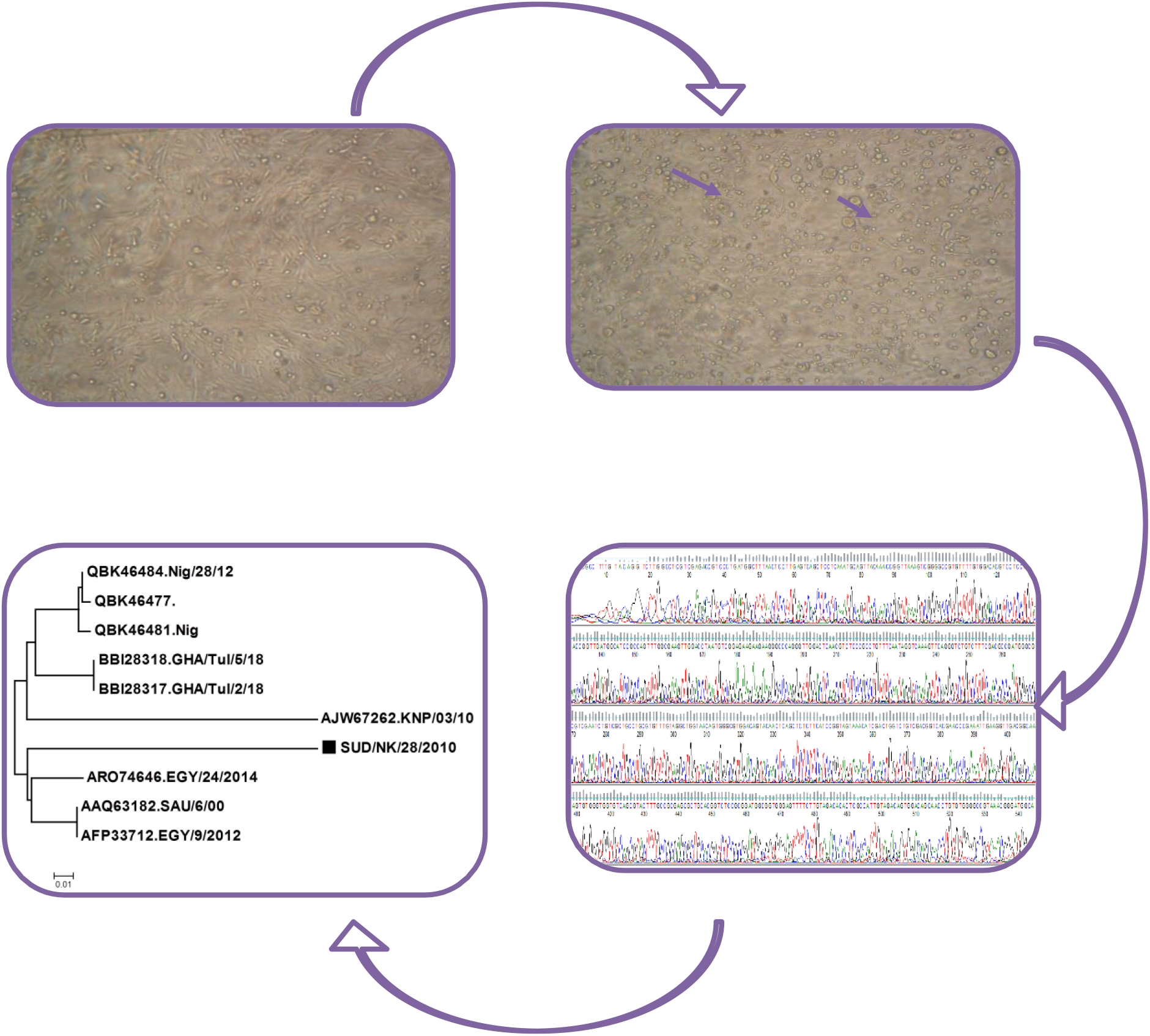
A: CPE developed by the virus showed rounding of cells pointed with black arrows was diffused in BHK cell line culture. B: Control BHK cell line clone 21. C: Nucleotides sequences of *SUD/NK/28/2010* poly protein histogram was visualized by Finich TV software. D: Dendogram of *FMD SAT2 SUD/NK/28/2010* serotype based on protein sequence of (P1) poly protein coding region showed that it is more related to Egyptian isolates (EGY/24/2014 and EGY/9/2012) and Saudi Arabia (SAU/6/00).

**Figure (2).**
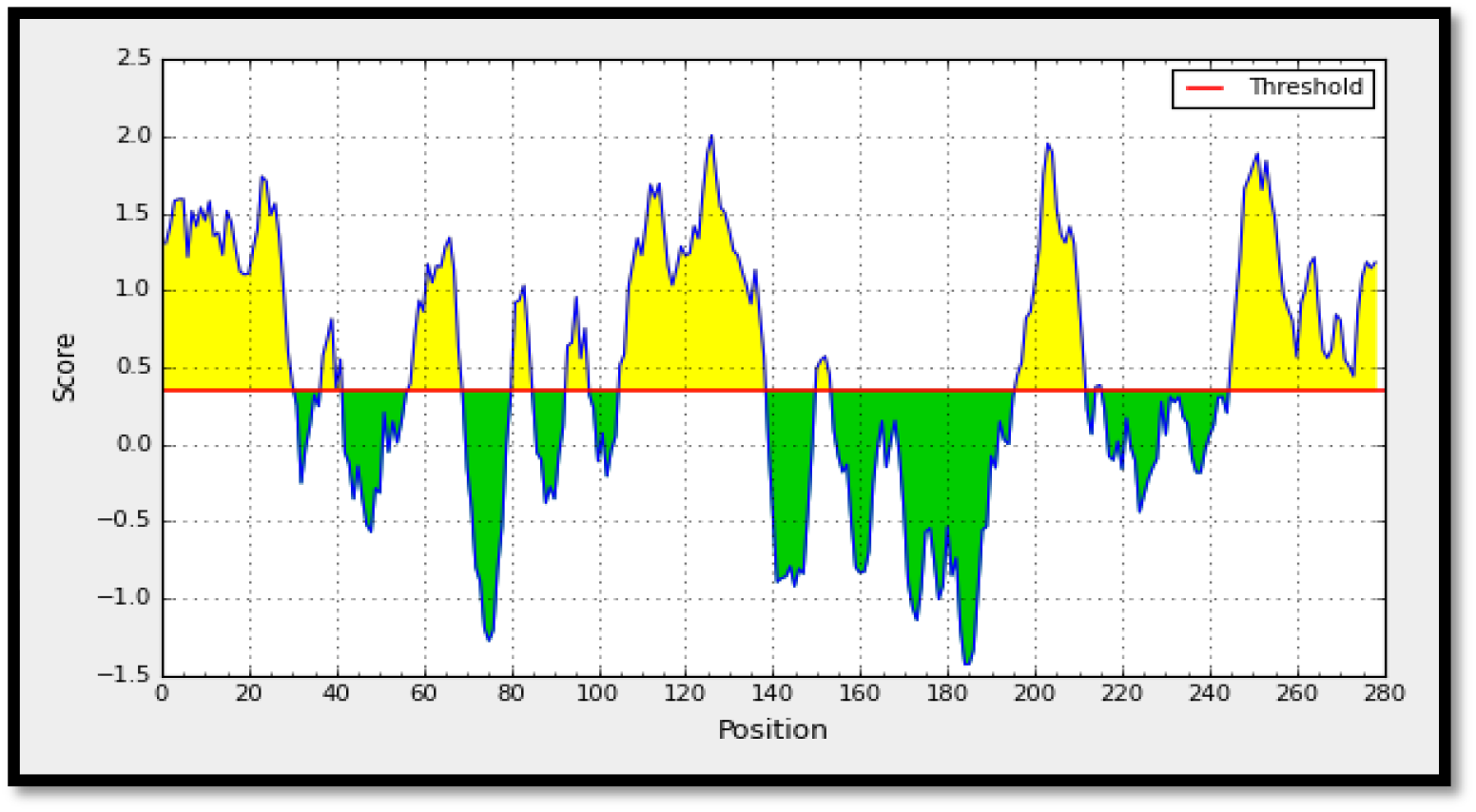
Present Bepipred Linear Epitope Prediction, the yellow space above threshold (red line) is proposed to be a part of B-cell epitopes and the green space is not a part.

**Figure (3).**
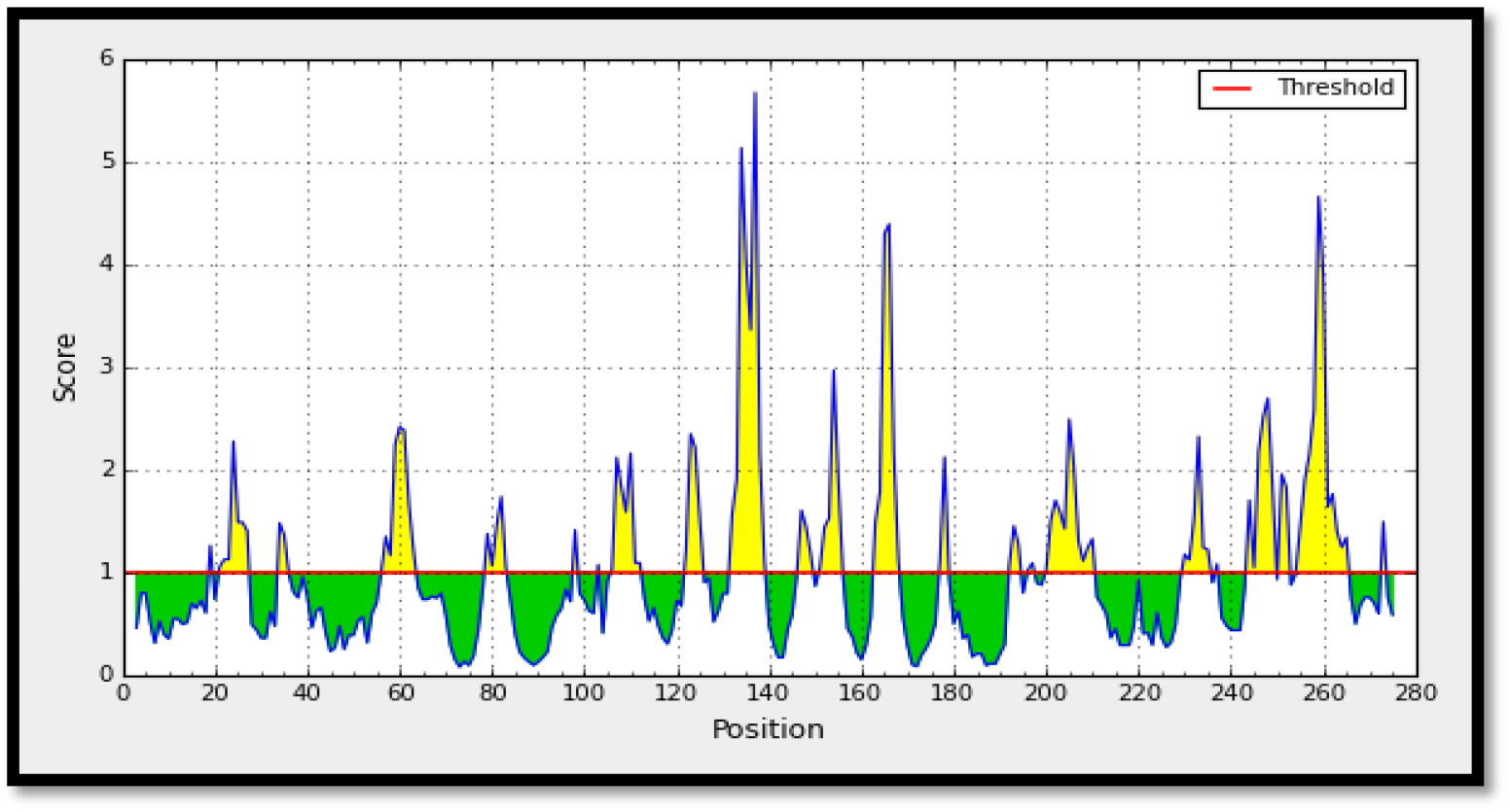
Emini surface accessibility prediction. The X-axes represent the sequence position and the Y-axes represent surface probability. The threshold value is 1.0 represented by the red line. The regions above the threshold are antigenic, shown in yellow and the region below the threshold is not antigenic, shown in green.

**Figure (4).**
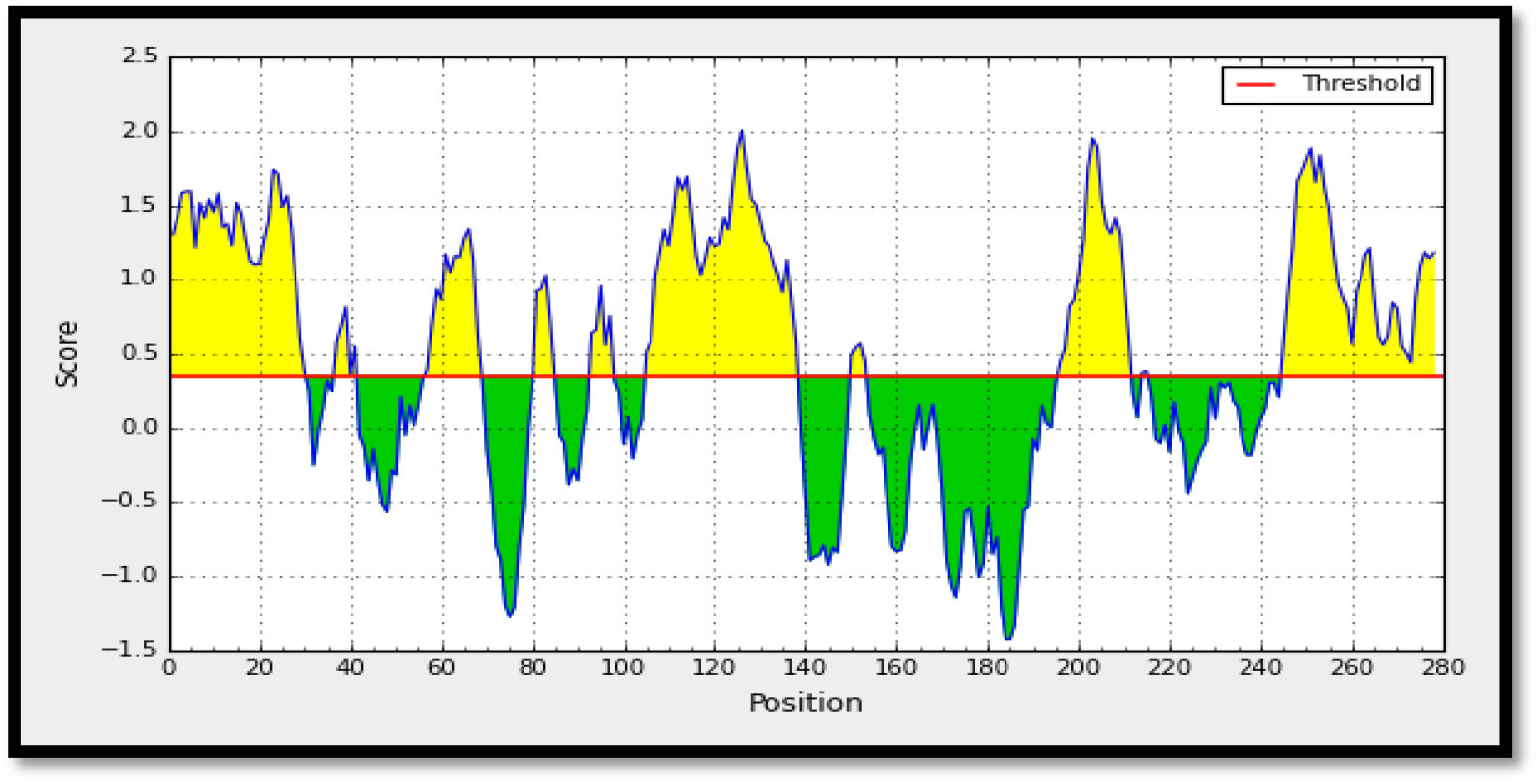
Kolaskar and Tongaonkar antigenicity prediction of the proposed epitope. Notes: The X-axes represent the sequence position and the Y-axes represent the antigenic propensity score. The threshold value is 1.0 represented by the red line. The regions above the threshold are antigenic, shown in yellow and the region below the threshold is not antigenic, shown in green.

### Prediction of T helper cell epitopes and interaction with MHC I alleles

Two promising peptides had binding affinity with MHC I and score IC50 ≤ 500. The lists of peptides were shown on table (2).

**Table (2):** List of the most promising T cell epitopes that had binding affinity with MHC I alleles along with their positions in the FMD SAT2, IC50, and percentile.

| Start | End | Epitope | Alleles | IC50 |
| --- | --- | --- | --- | --- |
| 256 | 264 | VQRSRQSTL | BoLA-D18.4 | 119.24 |
| 32 | 40 | YHAEWDTGL | BoLA-T2C | 473.49 |

### Physicochemical properties

The physicochemical properties of SUD/NK/28/2010 poly protein were assessed using BioEdit software version 7.0.9.0. The protein length was found to be 363 amino acids and the molecular weight was 39 675.83 Da. The amino acid that formed FMD SAT2 poly protein and their number along with their molar percentage (Mol%) were shown in Table 3 and Figure (5).

**Table (3):** Present the list of amino acid that formed SUD/NK/28/2010 protein with their number and Mol % using BioEdit software Version 7.2.6.

| Amino acid | number | MOI% | Amino acid | number | MOI% |
| --- | --- | --- | --- | --- | --- |
| Ala | 28 | 10.07 | Met M | 5 | 1.80 |
| Cys C | 4 | 1.44 | Asn N | 6 | 2.16 |
| Asp D | 17 | 6.12 | Pro P | 23 | 8.27 |
| Glu E | 9 | 3.24 | Gln Q | 10 | 3.60 |
| Phe F | 12 | 4.32 | Arg R | 15 | 5.40 |
| Gly G | 21 | 7.55 | Ser S | 21 | 7.55 |
| His H | 11 | 3.96 | Thr T | 33 | 11.87 |
| Ile I | 7 | 2.52 | Val V | 24 | 8.63 |
| Lys K | 7 | 2.52 | Trp W | 3 | 1.08 |
| Leu L | 17 | 6.12 | Try Y | 3 | 1.80 |

**Figure (5):**
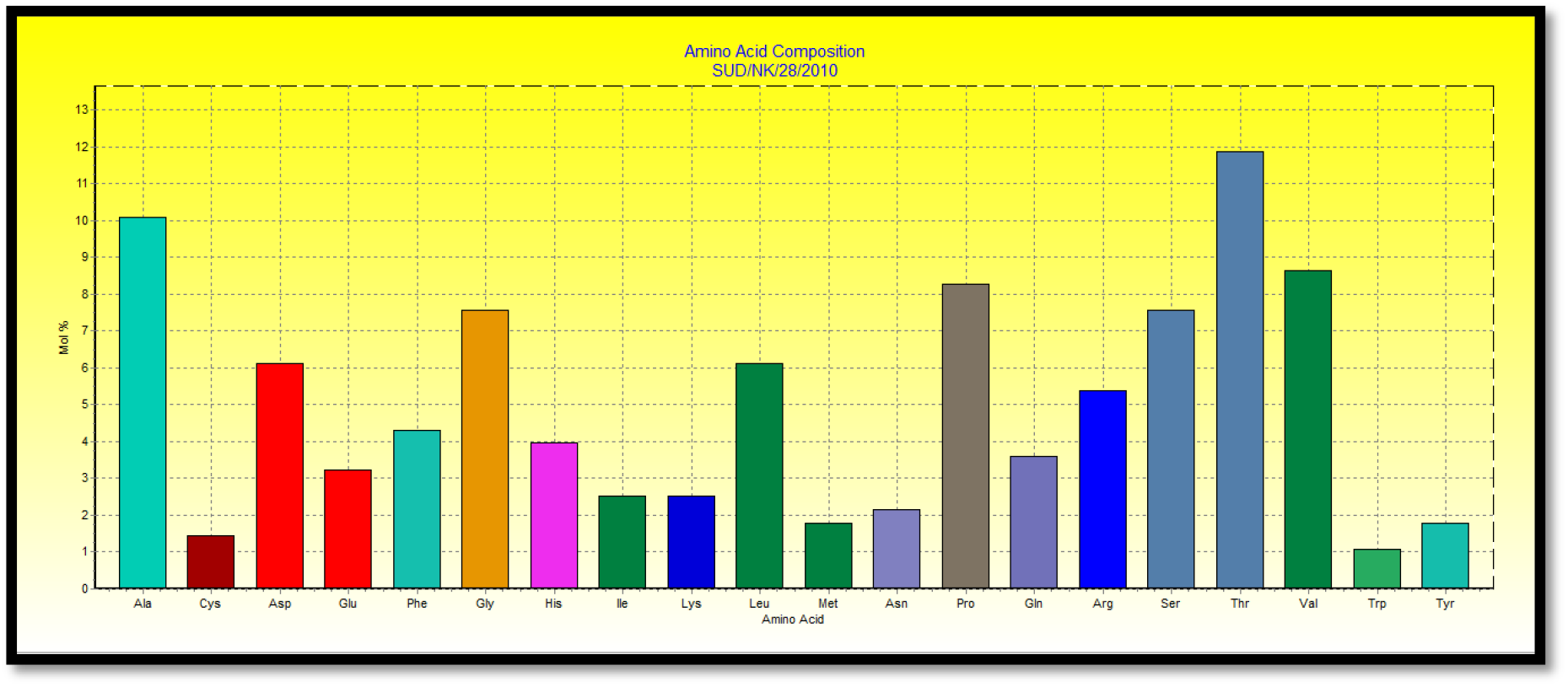
Amino acid composition in Mol % of *SAT2 SUD/NK/28/2010* using BioEdit software Version 7.2.6.

### Homology modeling

Two predicted T cell epitopes were found on VP1 protein. T cell epitope (*VQRSRQSTL*) started from 178 – 186 while T cell CD8 (*YHAEWDTGL*) started from 144 – 152. B cell epitope (*LPATPEDAAH*) was found on VP3 the epitope started from 133 – 141. All epitopes were demonstrated on 3 D structure of ref SAT2 poly protein (Maree *et al* 2011) figure (6).

**Figure (6):**
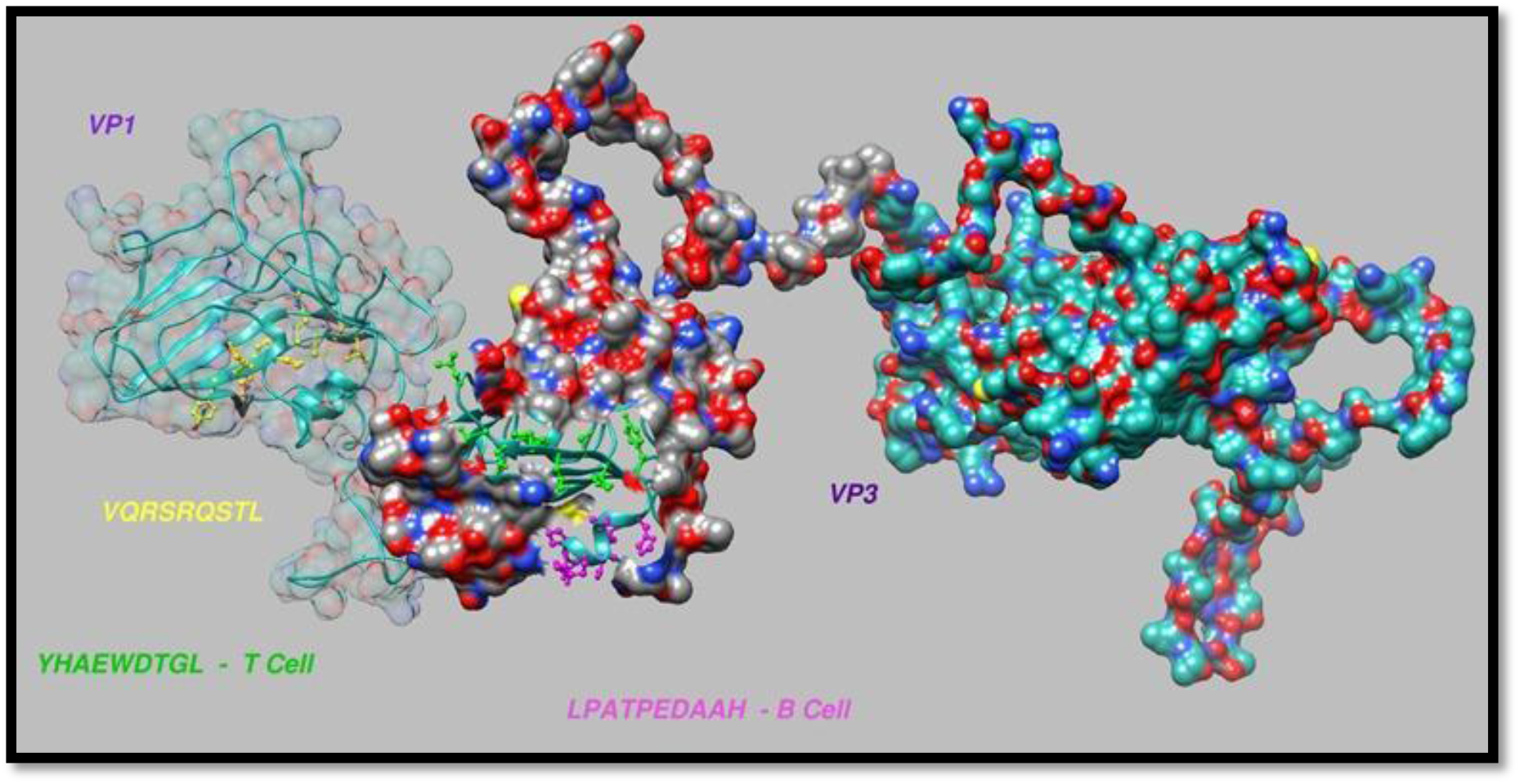
The 3D structure of ref SAT2 poly protein (Maree *et al* 2011) demonstrated T cell epitope CD8 (*VQRSRQSTL*) in VP1 protein (yellow color), T cell CD8 (*YHAEWDTGL*) (green color) and B cell (*LPATPEDAAH*) epitopes in VP3 protein (purple color).

### Molecular docking

Molecular docking was performed using (Moe 2007), T cell epitope (***VQRSRQSTL***) exhibits an exceptional result in term of strength of binding with ΔG value of value of - 26.24 kcal/mol when was docked with BoLA-HD6. The 3 D and 2 D structures were illustrated on figures (7 & 8). On the other hand, the core sequence **(*YHAEWDTGL***) had shown a less negative energy of binding in the binding site of BoLA-T2C (−23 kcal/mol), the 3 D and 2D structures were illustrated on figures (9 & 10).

**Figure (7):**
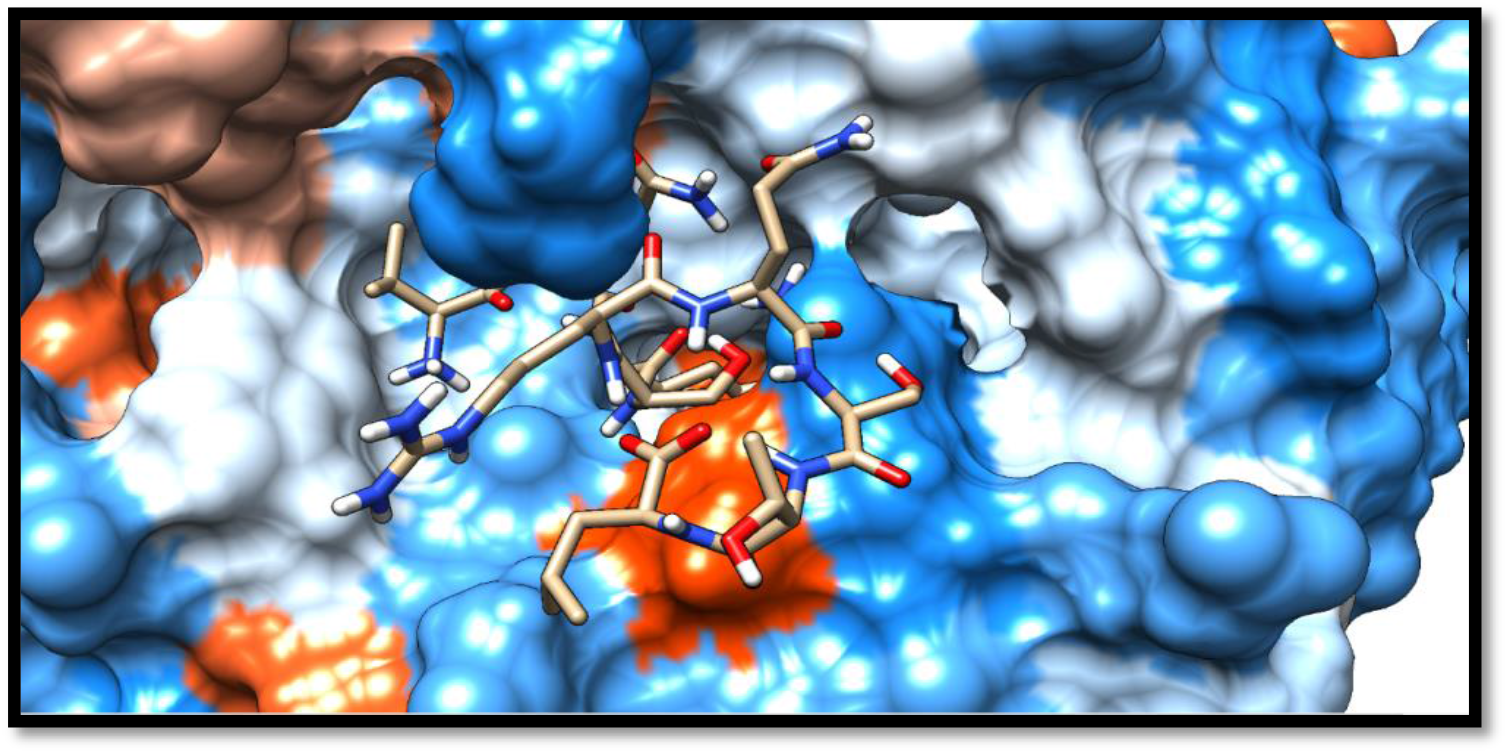
Illustrate the 3D interaction of the best docking pose of **(*VQRSRQSTL*)** in the binding sites of BoLA-HD6 using UCSF chimera 1.13.1 software.

**Figure (8):**
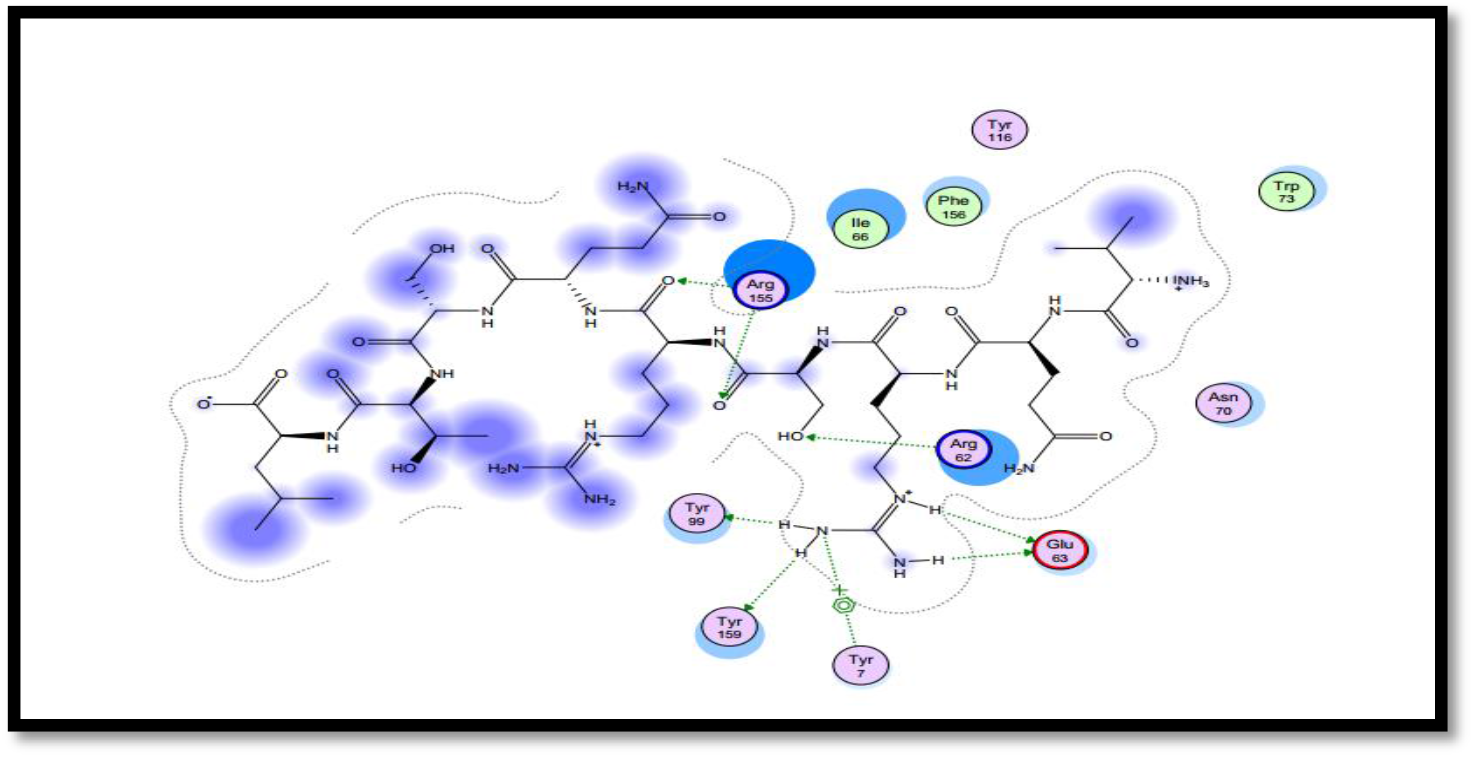
Illustrate the 2D interaction of the best docking pose of (***VQRSRQSTL)*** in the binding sites of BoLA-HD6 using MOE (2007).

**Figure (9):**
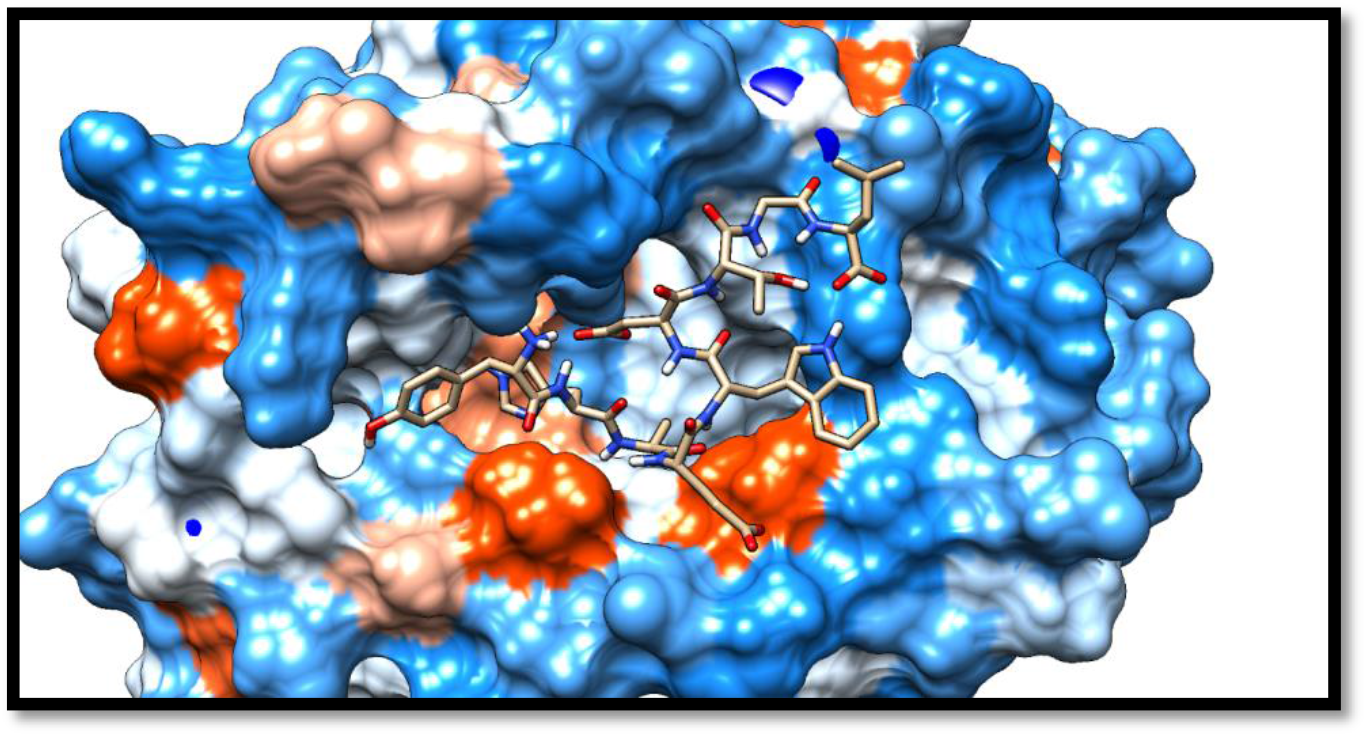
Illustrate the 3D interaction of the best docking pose of (***YHAEWDTGL***) in the binding sites of BoLA-T2C using UCSF chimera 1.13.1 software.

**Figure (10):**
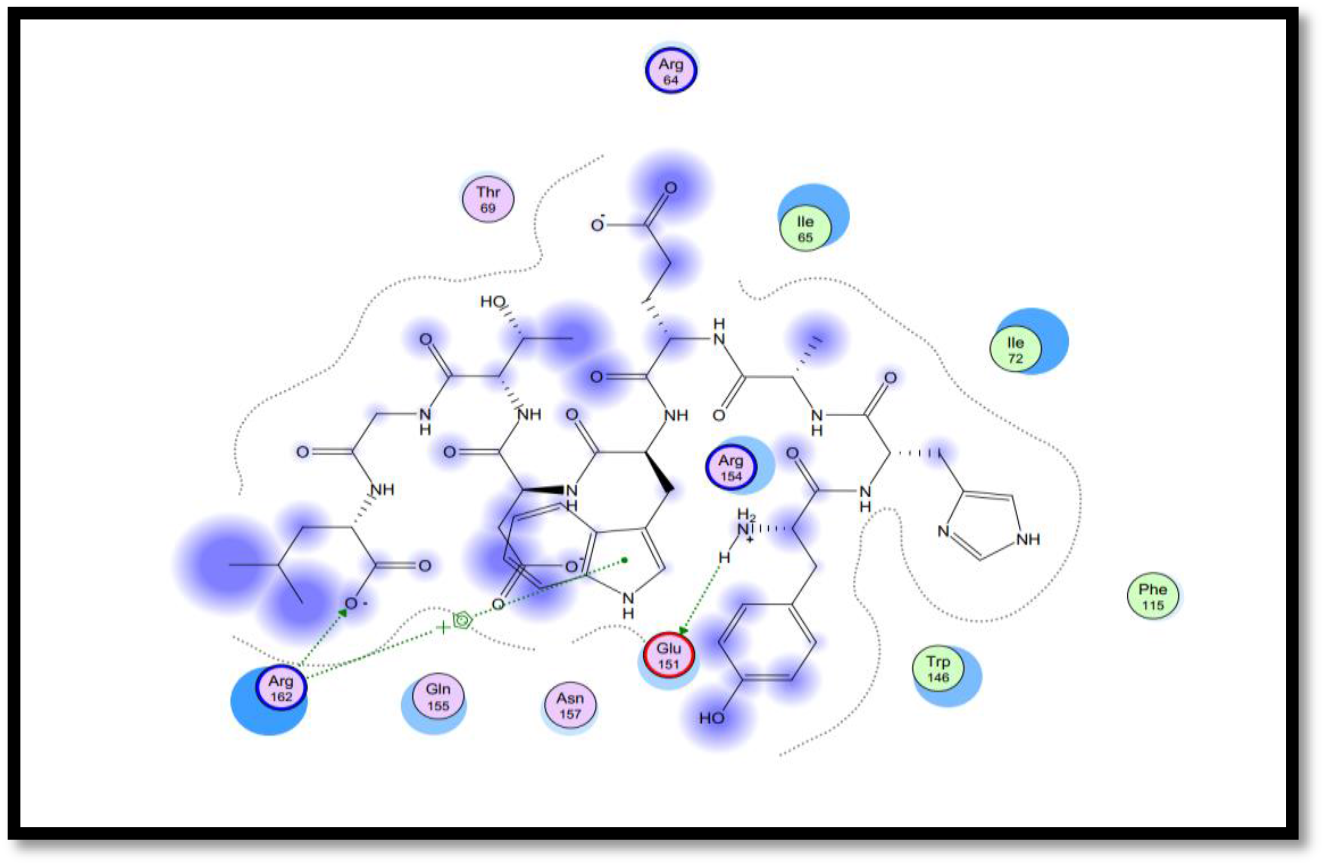
Illustrate the 2D interaction of the best docking pose of **(*YHAEWDTGL*)** in the binding sites of BoLA-T2C using MOE (2007).

## Discussion

Towards developing of FMD SAT2 epitope-based vaccine; *FMD/SAT2/NK/2010* virus was successfully adapted in BHK cell line as first step to develop vaccine (OIE 2017). Molecular characterization of FMD SAT2 was conducted using PCR with pair of primers that detect P1 gene region with expected amplicon size of 1279 bp in concordance with Knowles *et al* (2016). In the presented study, three novel epitopes were predicted in the Sudanese FMD SAT2 P1 polyprotein sequence; two of them were for T cells, and the third one was for B cells as a proposed candidate peptide vaccine. The T cell epitope (**VQRSRQSTL**) was located in VP1 protein in 178 –186 residues, while the other T cell epitope (**YHAEWDTGL**) and B cell epitope (**LPATPEDAAH**) were located in VP3 protein in 144–152 and 133–141 residues, respectively. Using ANN methods, the T cell epitopes (CD8) were predicted by analyzing the binding to the MHC1 molecule. High affinity epitopes that bind with BolA-D18.4 and BolA-T2C with an IC 50 of less than 500 acquire high affinity. According to Berzofsky *et al*. (2001) there are three ways for epitope enhancement. One of them is increasing the binding affinity to the major histocompatibility complex (MHC); this approach can effectively increase the potency of the vaccine (Berzofsky *et al*. 2001). MHC1 binding with the epitope is essential for an immune response that will produce CD8+ that is required for the elimination of viral infectivity (Groscurth and Filgueira 1998). MHC-I binding predictions have broad allelic coverage due to integration with proteasomal cleavage and TAP binding site predictions; 13 bovine leukocyte antigen class I (BolA-I) were described (Bamford et al. 1995; De Groot et al. 2003; Gaddum et al. 1996a; Gaddum et al. 1996b; MacHugh et al. 2011; Momtaz et al. 2014. MHC-II binding predictions are not as well developed as MHC-I binding predictions, yet they are still developing at a fast rate (Tong and Ren 2009; Momtaz *et al*. 2014; Al Asari et *al* 2017). In this study, we weren’t able to predict MHC-II epitopes in the virus protein due to a lack of alleles in the software special for bovine that would bind with the viruses that affected cattle. More efforts are required in this line of software development to eliminate this limitation. One B cell epitope (***LPATPEDAAH***) was predicted in the VP3 protein. B-cell epitopes are surface-accessible clusters of amino acids that can be identified by secreted antibodies or B-cell receptors and have the ability to induce cellular or humoral immune responses (Getzoff *et al*., 1988). The linear B cell epitopes of FMD P1 polyprotein were achieved by the Bepipred linear epitope prediction tool in (IEDB). Fully conserved epitopes were tested for surface accessibility and antigenicity with the aid of the Emini surface accessibility prediction method and the Kolaskar and Tongaonkar antigenicity methods, respectively; the peptides passed the default threshold value of 1.000 and 1.033, respectively.

According to their high scores in Emini surface accessibility and antigenicity tests, an epitope (***LPATPEDAAH***) was approved as a promising B cell epitope. The peptide passes the linearity test and this is important for a proposed B cell epitope for the vaccine; the immune system response is efficient with the linear peptide instead of a discontinuous epitope (Mauricio *et al*., 2004). Molecular docking was performed using (Moe 2007), T cell epitope (***VQRSRQSTL*)** exhibited an exceptional result in terms of strength of binding with ΔG value of-26.24 kcal/mol when docked with BoLA-HD6. On the other hand, the core sequence **(*YHAEWDTGL*)** has shown a lower negative energy of binding in the binding site of BoLA-T2C (−23 kcal/mol). This variation in docking score might be due to the number of amino acid residues that have some hydrophobic properties, which have a direct impact on the increase of binding energy during molecular docking (Meng et al. 2011), and it has been reported that the hydrophobic interaction may contribute as much as 50% of the total strength of the Ab-antigen bond (Virella 2001). It has been noticed for decades that FMDV has a high mutation rate and this leads to the rise of variants that might escape from the neutralizing activity of peptide antibodies that are directed against only one antigenic site. To overcome this limitation, a peptide vaccine should contain at least three or more different sites per virus type (Barteling and Woortmeyer 1987). Interestingly, the presented vaccine design contain three epitopes from different virus sites (VP1 and VP3), which might elicit different antibodies to neutralize the virus variants.

This study discovered some previously unknown predicted epitopes in Sudanese SAT2 topotype VII polyprotein. Many epitopes have been identified (Crowther et al 1993, Grazioli et al 2006, Opperman et al 2012, 2014) and predicted (Mukonyora 2015) in different regions of SAT2 capsid proteins in previous studies. However, no epitope has previously been observed in the same location as the epitope discovered in this study. This was not surprising given that SAT2 demonstrated high levels of genetic diversity in the P1-coding region within the SAT serotypes, with a 1.64 percent substitution rate observed on SAT2 per year (Vosloo et al 1996 and Phologane et al 2008).

This diversity has a direct impact on antigenicity and disease control through vaccination (Maree et al. 2011). In this regard, tailored diagnostics and vaccination are strongly advised for SAT2 control in the country. Nonstructural proteins that contain identified T cell epitopes have a strong correlation with antibodies that have a prolonged immune response (Barteling and Woortmeyer 1987; Blanco et al. 2001) and should be incorporated into vaccine designs.

To summarize, the preliminary results reported in this study necessitate testing the peptides after synthesis in target animals, as well as testing the potency of the vaccine in challenge animals after vaccination. These animal trials will allow calculating the PD50 of the vaccine and will be carried out in accordance with the recommended methods described for FMD vaccine (OIE 2017, Doel 1996, Cox *et al* 2005, Cox *et al* 2006, and Liu *et al* 2017).

### Study limitations

The poly protein sequence used in this study is a partial sequence of about 278 amino acids, which reflects in the number of predicted epitopes; however, the full genome sequence of the virus is highly recommended to predict new epitopes.

## Author Contributions

The analysis was conducted by: Inas, Ouabi, Isra, Sana, Ahmed, Mohamed, Mona, Sirdar. Interpretation of the analysis and writing of the manuscript were conducted by all authors. Supervision of the work was conducted by Dr. Mohamed Ahmed Salih, Dr. Reham Mohamed Elhassan, Yazeed Abdal Raouf and Muna Osman Alhaj

